# Genomes and transcriptomes help unravel the complex life cycle of the blastoclad fungus, *Coelomomyces lativittatus,* an obligate parasite of mosquitoes and microcrustaceans

**DOI:** 10.1101/2023.01.14.524055

**Authors:** Cassandra L. Ettinger, Talieh Ostovar, Mark Yacoub, Steven Ahrendt, Robert H. Hice, Brian A. Federici, Jason E. Stajich

## Abstract

Species of the phylum Blastocladiomycota, early diverging zoosporic (flagellated) lineages of fungi, are vastly understudied. This phylum includes the genus *Coelomomyces* which consists of more than 80 fungal species that are obligate parasites of arthropods. Known *Coelomomyces* species lack a complete asexual life cycle, instead surviving through an obligate heteroecious alternation of generations life cycle. Despite their global distribution and interesting life cycle, little is known about the genomics of any *Coelomomyces* species. To address this, we generated three draft-level genomes and annotations for *C. lativittatus* representing its haploid meiospore, orange gamete, and amber gamete life stages. These draft genome assemblies ranged in size from 5002 to 5799 contigs with a total length of 19.8-22.8 Mb and a mean of 7416 protein-coding genes. We then demonstrated the utility of these genomes by combining the draft annotations as a reference for analysis of *C. lativittatus* transcriptomes. We analyzed transcriptomes from across host-associated life stages including infection of larva and excised mature sporangia from the mosquito, *Anopheles quadrimaculatus*. We identified differentially expressed genes and enriched GO terms both across and within life stages and used these to make hypotheses about *C. lativittatus* biology. Generally, we found the *C. lativittatus* transcriptome to be a complex and dynamic expression landscape; GO terms related to metabolism and transport processes were enriched during infection and terms related to dispersal were enriched during sporulation. We further identified five HMG box genes in *C. lativittatus*, three belonging to clades with mating type (MAT) loci from other fungi, as well as four ortholog expansions in *C. lativittatus* compared to other fungi. The *C. lativittatus* genomes and transcriptomes reported here are a valuable resource and may be leveraged toward furthering understanding of the biology of these and other early diverging fungal lineages.

## INTRODUCTION

Fungi contribute to critical roles in the global ecosystem, yet knowledge of their biology, genetics and biochemistry largely stems from observations of only two phyla, the Ascomycota and Basidiomycota (i.e., the Dikarya). Zoosporic (flagellated) lineages of fungi make up additional fungal phyla (including the Blastocladiomycota and Chytridiomycota), but are generally understudied (James et al. 2020). The phylogenetic placement of these early-diverging zoosporic lineages is controversial and under constant revision as new genomic data becomes available (James et al. 2020; Li et al. 2021). For example, fungi belonging to the Blastocladiomycota were originally placed together with lineages in the Chytridiomycota, but now Blastocladiomycota is its own phylum (James et al. 2006; Porter et al. 2011). In addition, recent phylogenetic efforts have suggested that Blastocladiomycota may be more closely related to the Dikarya than the Chytridiomycota (Amses et al. 2022).

Within the Blastocladiomycota, the genus *Coelomomyces* (Blastocladiales; Coelomomycetaceae) consists of more than 80 highly fastidious fungal species that are obligate fatal parasites, primarily of mosquitoes and microcrustaceans (Whisler et al. 1975; Couch and Bland 1985; Powell 2017; Shen et al. 2020). These fungi have a worldwide distribution and over the last hundred years have been reported infecting all major genera of mosquitoes, i.e., *Aedes*, *Culex*, and *Anopheles*, each of which contain many species that transmit pathogens that cause medically important diseases such as malaria, filariasis, and various viral encephalitides. Moreover, because it is difficult to detect *Coelomomyces* infections in larval and adult mosquitoes, it is estimated that there are more than several hundred species worldwide yet to be described (Couch and Bland 1985), making these fungi a very large group for which we know virtually nothing about their genomes and biochemistry. This lack of knowledge is due to the failure, despite numerous attempts, to culture any species of *Coelomomyces in vitro*. One major difficulty is the lack of a cell wall on hyphae growing in the vegetative stages of their mosquito and copepod hosts. As far as is known, *Coelomomyces* species lack a complete asexual life cycle, instead surviving through an obligate alternation of generations in which a sporophytic phase parasitizes mosquitoes (e.g., larva) and a gametopytic phase parasitizes microcrustaceans (e.g., copepods) (Figure 1) (Whisler et al. 1975; Federici and Chapman 1977; Couch and Bland 1985). This type of lifecycle is uncommon in fungi, though a similar heteroecious life cycle is observed in the rust fungi (Duplessis et al. 2021).

**Figure 1.**
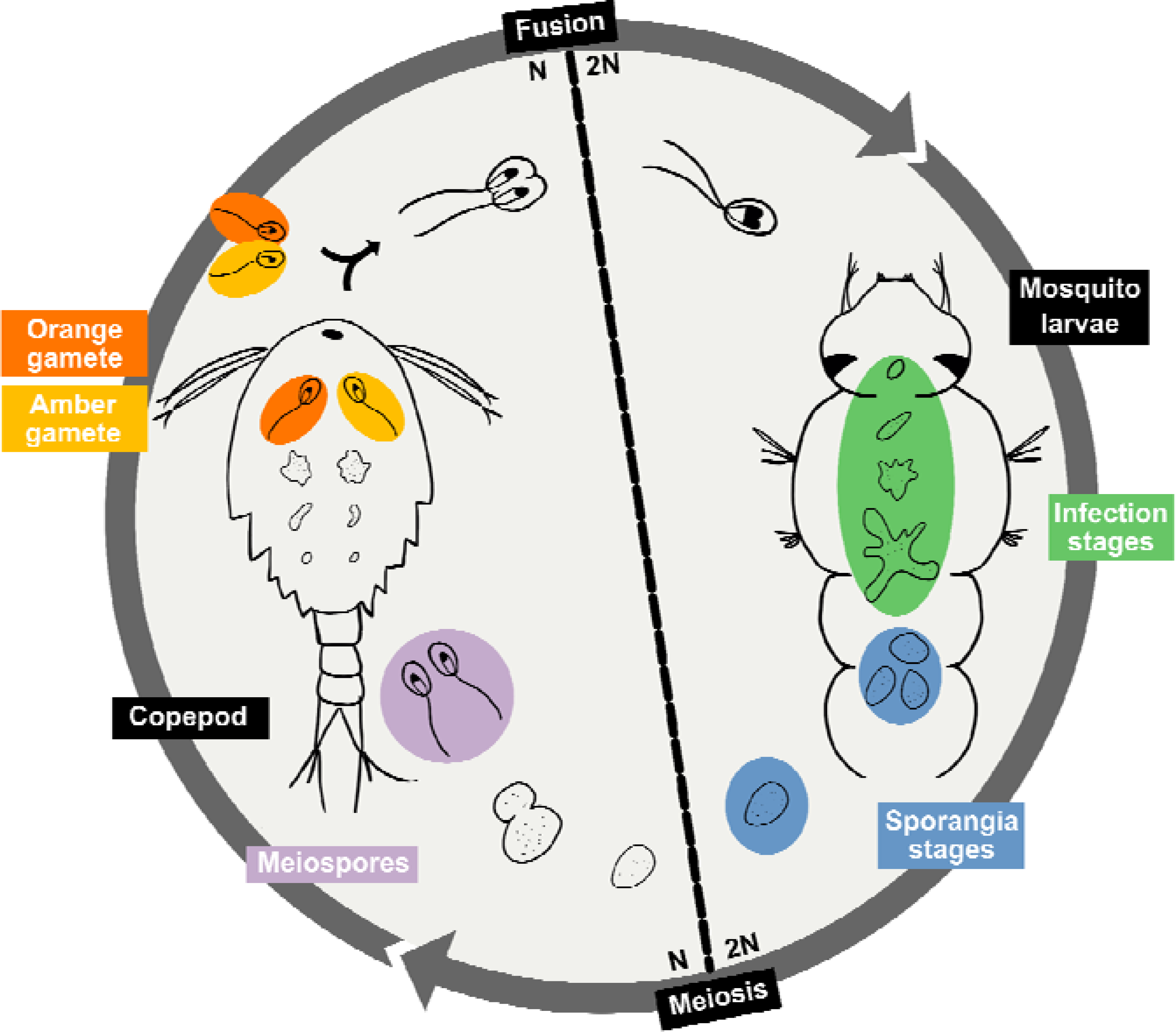
Alternation of generations life cycle of *Coelomomyces lativittatus*. Diagram showing the general alternation of generations life cycle of *C. lativittatus* between copepod and mosquito hosts. We have circled and highlighted the different life stages that were used in this study. Genomic sequencing was performed on haploid stages: including orange gametes (orange), amber gametes (yellow) and meiospores (purple). RNA sequencing was performed on diploid stages including mosquito larval infection stages (green) as well as sporangial stages excised from mosquito larva (blue).

The *Coelomomyces* life cycle begins when a biflagellate zygote encounters a mosquito larva. The motile spore encysts on the intersegmental membrane of the mosquito cuticle, a process facilitated by the secretion of adhesion vesicles (Travland 1979). The encysted spore develops an appressorium and penetration tube, which penetrates through the host cuticle (Zebold et al. 1979). Once inside the mosquito larva, the so-called hyphagens grow into coenocytic hyphae that ramify within the hemocoel over a period of seven to ten days, which then form sporangia at their tips (Federici and Chapman 1977; Couch and Bland 1985). The mosquito larva subsequently die and putrefy, liberating the sporangia. Meiosis then occurs in the sporangia resulting in haploid uniflagellate meiospores, which after sporangial dehiscence seek out and infect a crustacean host (typically copepods, though ostracods can serve as hosts as in some *Coelomomyces* species (Whisler et al. 2009)).

The penetration of copepods is thought to occur in a manner similar to that of the mosquito larva (Federici and Chapman 1977; Zebold et al. 1979), after which hyphae of the gametophyte form a holocarpic gametangium that cleaves into gametes. The meiospores that infect the copepods are of opposite mating types, thus forming gametangia in the copepod host that can generate gametes of opposite mating types. When gametogenesis is complete, the gametangium bursts, killing the copepod host and allowing the gametes to escape through fissures in the intersegmental membranes. If a copepod is infected by meiospores of each mating type, the gametangia burst simultaneously, and gametes of opposite mating types mate forming biflagellate zygotes within the dead copepod, which then seek out a mosquito larval host after release, thereby completing the alternations of generations life cycle (Whisler et al. 1975). If only a single mating type gametangium develops within a copepod, the gametes swim to the surface where they seek a mate in the water in which the mosquito larvae are breeding. In some species, such as *C. punctatus* and *C. dodgei*, the gametangia and gametes of different mating types, much like those of *Blastocladiella emersonii*, are different colors, apparently due to different levels of β-carotene, with one isoforms being bright orange, and the other light amber (Federici 1977; Federici and Thompson 1979).

As noted above, despite their worldwide distribution and relatively unique life cycle, little is known about the biology, biochemistry, or genomic landscape of *Coelomomyces* species. Modern molecular and genomic techniques allow us to circumvent the need for *in vitro* culturing and to expand foundational knowledge of this enigmatic fungal genus. Toward this goal, we have established an *in vivo* culture of *C. lativittatus*, a close relative of *C. dodgei* and *C. punctatus (Federici 1979; Couch and Bland 1985)*, which we maintain using the mosquito, *Anopheles quadrimaculatus*, and the copepod, *Acanthocyclops vernalis*. The research presented here represents the first exploratory investigation of *Coelomomyces* genomics and the *C. lativittatus* transcriptome.

To begin to answer questions related to *Coelomomyces* biology, we generated draft genomes and annotations for *C. lativittatus* from three life stages: (i) meiospores, (ii) orange gametes, and (iii) amber gametes. We generated transcriptomes from across life stages including infection of *An*. *quadrimaculatus* hosts and sporangia excised from *An. quadrimaculatus*, to elucidate genes involved in the unique biology and alternation of generations life cycle of this fungus. We then searched for mating type loci in *C. lativittatus*, as well as looked at expansions of orthologous genes compared to close relatives in the Blastocladiomycota and Chytridiomycota. The *C. lativittatus* genomes and transcriptomes reported here provide an invaluable foundational resource for understanding the biology of this and other unique and important understudied fungal lineages in various worldwide aquatic ecosystems.

## MATERIALS AND METHODS

### Study system

Larvae and copepods used for maintenance of the *in vivo* culture of *C. lativittatus* were, respectively, *Anopheles quadrimaculatus* and *Acanthocyclops vernalis*. These were maintained in culture as described previously (Federici 1983).

### DNA extraction methods & sequencing

We sought to generate genomes from three haploid life stages (i) meiospores, (ii) orange gametes, and (iii) amber gametes. To obtain *C. lativittatus* meiospores, mosquito larvae with advanced infections were collected when full of sporangia, within a day or two of death. To induce germination of sporangia, each larva was surface-sterilized by rinsing it in 70% ethanol for 20 seconds, after which each was placed in 1 mL of double-distilled water in a 22 mm plastic Petri dish at room temperature. The larvae were dissected with jeweler’s forceps and most of the cuticle and midgut were removed from the water. Typically, the sporangia dehisced, releasing meiospores, 48-72 hours after incubation at room temperature. Meiospore samples were collected using a 1 mL pipette and centrifuged using a table-top mini-fuge for 3 seconds to sediment any sporangia in the sample. The meiospores were then pelleted by centrifugation at 16,000x*g* for 2 minutes. To obtain *C. lativittatus* gametes, infected copepods containing the orange and amber mating types were separated prior to copepod lysis. Liquid was removed and then copepods were surface-sterilized by rinsing in 70% ethanol and then washed with double-distilled water to reduce contaminants. After the gametes were released from the copepods, the copepod carcasses were removed by allowing them to settle in the microfuge tube and the supernatant containing the gametes was transferred to a new tube. Samples were spun for 3 minutes at 6000x*g* to pellet the gametes, and the supernatant was removed.

DNA was extracted from the resulting meiospore and gamete pellets using a Qiagen genomic DNA purification kit with Qiagen genomic-20/G tips following the standard manufacturer protocol. DNA was then amplified using the Qiagen repli-G Whole Genome Amplification Kit according to the standard manufacturer protocol. Illumina Libraries were prepared with the NEB DNA Library Prep Kit. Libraries were sequenced on an Illumina HiSeq 2500 with 100 bp PE sequencing by the UC Riverside Genomics Core Facilities.

### RNA extraction and library preparation

For transcriptome analysis of the sporophyte, hyphae were excised from infected fourth instar larvae of *An. quadrimaculatus* during either early, middle, or late stages of fungal development. For the purpose of this study, we define early, middle, and late infection stages as follows. Typically, the early stage of obvious infection appears as a few unpigmented, i.e., white, sporangia at the tips of hyphae about six days after molting to the fourth instar. The fat body in these larvae is quite well developed, and hyphae can be seen adhering to this tissue in each of the larval abdominal segments and in the thorax. The middle stage occurs over days seven and eight, during which the number of sporangia increases significantly, with most being mature, meaning rusty brown in color. The late stage occurs during days nine and ten, by which time many larvae are full of sporangia and die, although many other larvae survive another four to five days before dying. These larvae continue to grow, being at least twice the size at which healthy larvae pupate. Two replicate samples were collected from each time point.

For the sporangia transcriptomes, mosquito larvae with advanced infections were collected when full of sporangia, within a day or two of death. To induce sporangia to undergo meiosis and germinate, each larva was surface-sterilized by rinsing it in 70% ethanol for 20 seconds, after which each was placed in 1 mL of double-distilled water in a 22 mm plastic Petri dish at room temperature. The sporangia were excised from the larvae using jeweler’s forceps, after which the cuticle and midgut were removed from the water. Typically, the sporangia dehisced, releasing meiospores, 48-72 hours after incubation at room temperature. Sporangia were collected at 0, 24, 36, and 48 hour time points starting from the period the sporangia were excised from the mosquito larva (0 hour) through dehiscence (48 hour), when the uniflagellate meiospores burst out of the sporangia. Samples were collected using a 1 mL pipette and centrifuged using a table-top mini-fuge for 3 seconds to obtain the sporangia in the sample. Two replicate samples were collected from each time point.

RNA from all samples was extracted with Trizol (Life Technologies, Grand Island, NY) as per the manufacturer protocol. 1.2 μg of RNA was used as the starting material for the NEBNext Ultra Directional RNA Library Prep Kit for Illumina (New England BioLabs, Ipswich, MA). Poly-A RNA was purified as per instructions and converted to adapter-ligated, size-selected cDNA. An aliquot of the library was cloned into pJet1.2 (Thermo Fisher Scientific, Waltham, MA) and clones sequenced with standard methods to check library quality. An aliquot was also run on a Bioanalyzer 2100 (Agilent Technologies, Santa Clara, CA). The resulting sequencing libraries were sequenced by the Institute for Integrative Genome Biology Core facility at the University of California at Riverside using the MiSeq instrument with 100 bp paired-end reads (Illumina, San Diego, CA).

### Genome assembly

Genomes for the meiospore (MEIOSPORE), orange gamete (ORANGE) and amber gamete (AMBER) libraries were assembled using the Automatic Assembly of the Fungi (AAFTF) pipeline v. 0.2.3 (Stajich and Palmer 2019). Briefly, this involved first trimming and filtering reads using bbduk.sh from BBTools v. 37.76 (Bushnell 2014). Next, assemblies were produced using the ‘assemble’ command in AAFTF which relies on SPAdes v. 3.12.0 (Prjibelski et al. 2020) run with default parameters to select optimal *k*-mer size and screened for contaminant vectors with AAFTF vecscreen step using NCBI BLAST (Camacho et al. 2009). The AAFTF sourpurge step was run which utilizes sourmash v. 3.5.0 (Brown et al. 2016) to further purge any remaining contaminant contigs and AAFTF rmdup step using Minimap2 v. 2.17 (Li 2018) was run to identify duplicate contigs for removal. Finally, AAFTF polish step ran Pilon v. 1.22 (Walker et al. 2014) to polish the resulting contigs in each assembly to remove potentially mis-called consensus nucleotides or indels by SPAdes.

Assembly evaluation for each genome was performed using QUAST v. 5.0.0 (Gurevich et al. 2013) and BUSCO v. 5.0.0 (Simão et al. 2015) against both the eukaryote_odb10 and fungi_odb10 gene sets. BUSCO assessment was also performed for reference genomes from other fungal lineages for comparison (for lineages see Table S1). We performed telomere searches against the *Coelomomyces* assemblies using find_telomeres.py (https://github.com/markhilt/genome_analysis_tools) to test for telomeric repeats at the ends of the scaffolds and determine chromosome completeness (Hiltunen et al. 2021).

### Contamination screen and removal

Given the obligate nature of *C. lativittaus* with its hosts, microbial contamination was assessed in each assembly using the BlobTools2 pipeline (Figure S1) (Challis et al. 2020). This involved first assigning taxonomy against the UniProt Reference Proteomes database (v. 2020_10) to each contig using diamond (v. 2.0.4) and command-line BLAST v. 2.2.30+ (Camacho et al. 2009; Buchfink et al. 2021). Next, read coverage was calculated by aligning the reads from the MEIOSPORE, AMBER and ORANGE libraries to their respective genome assemblies with BWA v.0.7.17 (Li and Durbin 2009) and sorted using samtools v. 1.11 (Li et al. 2009). Finally, the BlobToolKit Viewer was used to visualize the GC content, read coverage, and predicted taxonomies of contigs to identify contaminants. This approach flagged 1969, 11, and 24 contigs as putative contaminants in the AMBER, MEIOSPORE and ORANGE assemblies respectively.

Microbial contamination was further assessed with the anvi’o v.7 pipeline (Eren et al. 2015, 2021), a complementary method, for the AMBER assembly given the large number of contaminants predicted by BlobTools2 (Figure S1). This involved first obtaining read coverage from each of the three genomic samples (AMBER, ORANGE, and MEIOSPORE) to the AMBER assembly with bowtie2 v. 2.4.2 (Langmead et al. 2009) and samtools v. 1.11 (Li et al. 2009). A contig database for the AMBER assembly was then generated using ‘anvi-gen-contigs-database’ which uses Prodigal v. 2.6.3 (Hyatt et al. 2010) to predict open-reading frames. This command also identifies single-copy bacterial (Lee 2019), archaeal (Lee 2019) and protista (Delmont 2018) genes using HMMER v. 3.2.1 (Eddy 2011) and ribosomal RNA genes using barrnap (Seemann 2018). We predicted taxonomy for each predicted ORF using Kaiju v. 1.7.2 (Menzel et al. 2016) with the NCBI BLAST non-redundant protein database nr including fungi and microbial eukaryotes v. 2020-05-25. We then constructed anvi’o profiles for each sample (AMBER, ORANGE, and MEIOSPORE) using ‘anvi-profile’ with the ‘--cluster-contigs’ option and a contig length cut-off of >2.5 kbp. These profiles were then merged using ‘anvi-merge’. Contaminant contigs in the AMBER assembly were then identified through ‘anvi-interactive’ using a combination of hierarchical clustering based on coverage and tetranucleotide frequency, taxonomic identity, and GC content. This second method identified 1127 contaminant contigs in the AMBER assembly.

Contaminant contigs (e.g., any contig identified by the BlobTools2 pipeline as assigned to bacteria, archaea or viruses and any contig identified using the anvi’o pipeline) were subsequently removed from the draft assemblies. For the AMBER assembly, this meant conservatively removing a total of 2091 contaminant contigs (1005 identified by both methods, 964 contigs identified by BlobTool2 only, and 122 identified by anvi’o only).

### Genome annotation

Genome annotation was performed using the Funannotate pipeline v.1.7.4 (Palmer and Stajich 2020). This first involved using RepeatModeler v.1.0.11 (Flynn et al. 2020) and RepeatMasker v.4.0.7 (Smit et al. 2013-2015) to generate a library of predicted repetitive elements and then soft mask these elements in the draft genomes. Next the RNASeq data was assembled using Trinity v. 2.10.0 in Genome-Guided mode and aligned with PASA v.2.3.3 to train the *ab initio* gene prediction algorithms augustus and SNAP (Haas et al. 2003; Grabherr et al. 2011). Consensus gene models were generated using EVidenceModeler v.1.1.1 (Haas et al. 2008) on predicted protein-coding gene models from a combination of algorithms including CodingQuarry v. 2.0, Augustus v. 3.3.3, GeneMark-ETS v. 4.38, GlimmerHMM, and SNAP v 2013_11_29 (Korf 2004; Majoros et al. 2004; Stanke et al. 2006; Ter-Hovhannisyan et al. 2008; Testa et al. 2015). Transfer rRNA genes were predicted using tRNAscan-SE v. 1.3.1 (Lowe and Eddy 1997). Protein annotations were predicted for the consensus gene models based on similarity to Pfam (Finn et al. 2014), CAZyme domains (Lombard et al. 2014; Huang et al. 2018), MEROPS (Rawlings et al. 2014), eggNOG v. 1.0.3 (Huerta-Cepas et al. 2016), InterProScan v. 5 (Jones et al. 2014), and Swiss-Prot (Boutet et al. 2016) using HMMER v.3 (Eddy 2011) or diamond BLASTP (Buchfink et al. 2015). Phobius (Käll et al. 2004) and SignalP v. 4.1c (Armenteros et al. 2019) were also run to predict transmembrane proteins and secreted proteins respectively. Any problematic gene models that were flagged by Funannotate were manually curated as needed. The annotation results were summarized in custom code written in R v. 4.0.3 using the tidyverse v. 1.3.0 package (Wickham et al. 2019; R Core Team 2020). The annotated genomes of the MEIOSPORE, ORANGE, and AMBER assemblies were aligned and mapped to the RNA sequencing reads using HISAT2 (Kim et al. 2019).

### Transcriptome analysis

Given the obligate relationship of *Coelomomyces* with its hosts, we chose a reference-based transcriptome approach as initial results from *de-novo* approaches revealed host contamination even after removal using a reference host transcriptome. To provide a more comprehensive gene set to use as a reference for transcriptome analysis, we combined the predicted transcript sets from all three genome annotations. We then used CD-HIT-EST to cluster transcripts at 90% sequence identity and evaluated this combined set (AOM90) with BUSCO v. 5.0.0 (Fu et al. 2012; Simão et al. 2015). For comparative purposes, a protein alignment of *C. lativittatus* with other fungal lineages (for lineages see Table S1), was constructed using the PHYling_unified (https://github.com/stajichlab/PHYling_unified) pipeline, which uses HMMER v.3 (Eddy 2011) and ClipKIT (Steenwyk et al. 2020) to search for markers in the protein sequences, build, and trim an alignment based on BUSCO fungi_odb10 HMMs. A maximum likelihood phylogeny was built from this alignment using IQ-TREE2 v.2.2.0 (Minh et al. 2020). BUSCO fungi_odb10 gene partitions were provided to IQ-TREE2 using -p (Chernomor et al. 2016) and ModelFinder Plus was run using -m MFP to ensure use of the best evolutionary model for each partition based on BIC (Kalyaanamoorthy et al. 2017).

Mosquito host transcripts were removed from the transcriptome data using BBMap against an *An. quadrimaculatus* (Accession: GBTE00000000) reference transcriptome prior to read quantification (Bushnell 2014; Desjardins et al. 2015). Host-filtered transcriptome read counts were quantified against the AOM90 transcript set using kallisto (Bray et al. 2016). The count data were then imported into R for analysis with the DESeq2, ggplot2 and GSEABase packages (Wickham 2009; Love et al. 2014; R Core Team 2020; Morgan et al. 2022).

General expression trends across all samples were visualized using variance stabilized count data. We then used DESeq2 to identify differentially expressed transcripts between life stages (e.g., sporangial vs. infection). Significant genes were defined as transcripts with a false discovery rate adjusted *p*-value of < 0.01 and a |log2 fold change| > 2. Gene Ontology (GO) enrichment analysis was performed to assess if the differentially expressed transcripts were significantly enriched in any particular functions (*p* < 0.05). This analysis was performed at each of three GO classes: biological processes (BP), molecular functions (MF), and cellular components (CC). Significantly enriched GO terms were simplified using Revigo with the default settings (Supek et al. 2011).

In order to identify transcripts that show change in expression across the development time course conditions within each sporangium (e.g., 0 hr vs. 24 hr vs. 32 hr vs. 48 hr) and infection stages (e.g., early vs. middle vs. late infection), we performed a Likelihood Ratio Test (LRT). Significant transcripts from LRT were filtered with a false discovery rate adjusted *p*-value of < 0.01 and a |log2 fold change| > 2. The “DEGreport” package was used to cluster genes with similar expression profiles based on the LRT results across different time series conditions (Pantano 2022). A GO enrichment analysis was performed to identify enriched GO terms in each of the clusters with similar expression patterns (*p* < 0.05). GO terms were simplified using Revigo with default settings (Supek et al. 2011).

### Identification of mating type (MAT) loci in C. lativattus

To identify high mobility group box (HMG-box) genes putatively involved in mating in the *C. lativittatus* genomes, we used HMMsearch v. 3.3.2 for PFAM PF00505 with an e-value of 1e-15 (Eddy 2011). Given that genes neighboring HMG-boxes are thought to be involved in mating in other fungi (Vossenberg et al. 2019), we used Clinker (Gilchrist and Chooi 2021) and Cblaster (Gilchrist et al. 2021) to assess the syntenic regions surrounding the HMG-boxes in the *Coelomomyces* assemblies to identify conserved regions neighboring HMG-boxes. To confirm phylogenetic placement of the identified HMG-box genes, we aligned the candidate genes from *C. lativattatus* with those of other fungi (for fungi used see Table S1) using HMMalign (Eddy 2011). We then constructed a maximum likelihood phylogenetic tree of the HMG-box genes using IQ-TREE using the VT+R6 model which was selected by ModelFinder Plus (Minh et al. 2020). Finally, to compare expression of HMG-box genes across *C. lativittatus* life stages, the variance stabilized expression levels of the HMG-box genes were plotted using ggplot2 (Wickham 2009). Pairwise t-tests were performed to assess differential expression between life stages.

### Identification of orthologous gene expansions relative to other fungal lineages

Orthofinder v. 2.5.4 was used to identify whether any differentially expressed transcripts represented genes expanded in copy number in *C. lativittatus* compared to other fungi (for fungi used see Table S1) (Emms and Kelly 2019). We filtered the Orthofinder results to orthogroups containing genes with differentially expressed transcripts in the RNAseq data. These results were subsetted by orthogroups that were at least |log2 fold change| > 2 higher in copy number in *C. lativittatus* compared to the other fungi. Orthogroup expansions were confirmed through phylogenetic analyses. Briefly, a nucleotide alignment of all genes in an orthogroup of interest was produced using MUSCLE v. 5.1 and then a phylogenetic tree was built with IQ-TREE2 using -m MFP which runs ModelFinder Plus (Kalyaanamoorthy et al. 2017).

## RESULTS

*Over half of the genomic landscape is represented in C. lativittatus assemblies and annotations* After successful contaminant removal, the draft genome assemblies ranged in size from 5002 to 5799 contigs with a total length between 19.8 Mb and 22.8 Mb (Table 1). Although the assemblies were fragmented with an average N50 of 6128 bp, BUSCO assessment found that the draft assemblies were halfway complete (Table S2). Mean completeness in ‘genome’ mode was 43.1% and 56.3% using the fungi_odb10 and eukaryota_odb10 sets, respectively. While, these values were lower than those of other blastoclads on average (fungi_odb10: 75.5%, eukaryota_odb10: 83.8%), including a recent long-read genome from *B. emersonii* (fungi_odb10: 81.8%) (Leonard et al. 2022), it is important to note that BUSCO sets are biased against early diverging fungal lineages. Nonetheless, these draft assemblies provide a valuable starting point for further improvement and refinement moving forward.

**Table 1.** Genome assembly and annotation statistics. Various statistics calculated by QUAST for each of the assemblies are provided here including the total number of contigs in the assembly, the total assembly length, percent GC content, the N50 and the L50. All statistics from QUAST are based on contigs of size >= 500 bp, unless specifically noted (e.g., “# contigs (>= 0 bp)” and “Total length (>= 0 bp)” include all contigs in each assembly). We also report the percentage of RNAseq reads that were aligned to each assembly using HISAT, the percentage of each assembly masked by RepeatMasker (this value includes total interspersed repeats, simple repeats and low complexity regions), and the number of predicted telomeres. A summary of genome annotation results is reported here including the total number of gene models, mRNAs, tRNAs, the mean exon length (bp), tRNA length (bp), mRNA length (bp), and the percentage of proteins with a PFAM domain, InterProScan, EggNog, COG, or GO term matches.

| Software | Statistic | Amber | Orange | Meiospore |
| --- | --- | --- | --- | --- |
| QUAST | # contigs ($\geq 0$ bp) | 5799 | 5413 | 5002 |
|  | Largest contig | 56469 | 30249 | 37025 |
| | Total length ($\geq 0$ bp) | 22882397 | 19891335 | 21857322 |
|  | GC (%) | 33.43 | 32.32 | 32.33 |
|  | N50 | 6067 | 5623 | 6695 |
|  | N90 | 1782 | 1603 | 1996 |
|  | L50 | 1089 | 1030 | 988 |
|  | L90 | 3884 | 3610 | 3254 |
|  | # N's per 100 kbp | 480.4 | 362.27 | 101.38 |
| **Telomeres** | Telomeres TOTAL | 23 | 18 | 22 |
|  | Telomeres FRWRD | 8 | 9 | 10 |
|  | Telomeres RVRS | 15 | 9 | 12 |
| **HISAT2** | Overall RNAseq alignment to genome | 55.40% | 55.17% | 59.01% |
| **RepeatMasker** | Repetitive regions (%) | 10.77% | 12.57% | 14.04% |
| **Funannotate** | Total number of gene models | 7404 | 7344 | 7677 |
|  | Number of mRNA genes | 7352 | 7289 | 7608 |
|  | Number of tRNA genes | 52 | 55 | 69 |
|  | Mean exon length | 367.89 | 358.18 | 372.63 |
|  | Mean tRNA length | 74.58 | 74.62 | 74.91 |
|  | Mean mRNA length | 1501.03 | 1309.02 | 1465.69 |
|  | Genes with PFAM hit (%) | 48.74 | 44.91 | 48.04 |
|  | Genes with InterProScan hit (%) | 65.11 | 63.82 | 64.91 |
|  | Genes with EggNog hit (%) | 54.16 | 52.78 | 53.82 |
|  | Genes with COG hit (%) | 53.28 | 52.03 | 52.98 |
|  | Genes with GO Term (%) | 46.83 | 45.68 | 46.72 |

Annotation of the three assemblies with Funannotate identified on average 7416 protein-coding genes and 59 tRNA genes, with 63.2% of these having a hit to at least one functional database (Table 1). We combined the three predicted transcript sets together at 90% identity using CD-HIT-EST to generate a comprehensive final gene set (AOM90) of 8645 transcripts and leading to improved BUSCO ‘protein’ scores of 62.5% and 82.8% using the fungi_odb10 and eukaryota_odb10 sets, respectively (Table S2; Figure S2). Despite being slightly lower than the average scores for other blastoclads (Figure 2; fungi_odb10: 84.1, eukaryota_odb10: 90.1%), the AOM90 transcript set represents a promising and robust reference for beginning to elucidate *C. lativittatus* biology.

**Figure 2.**
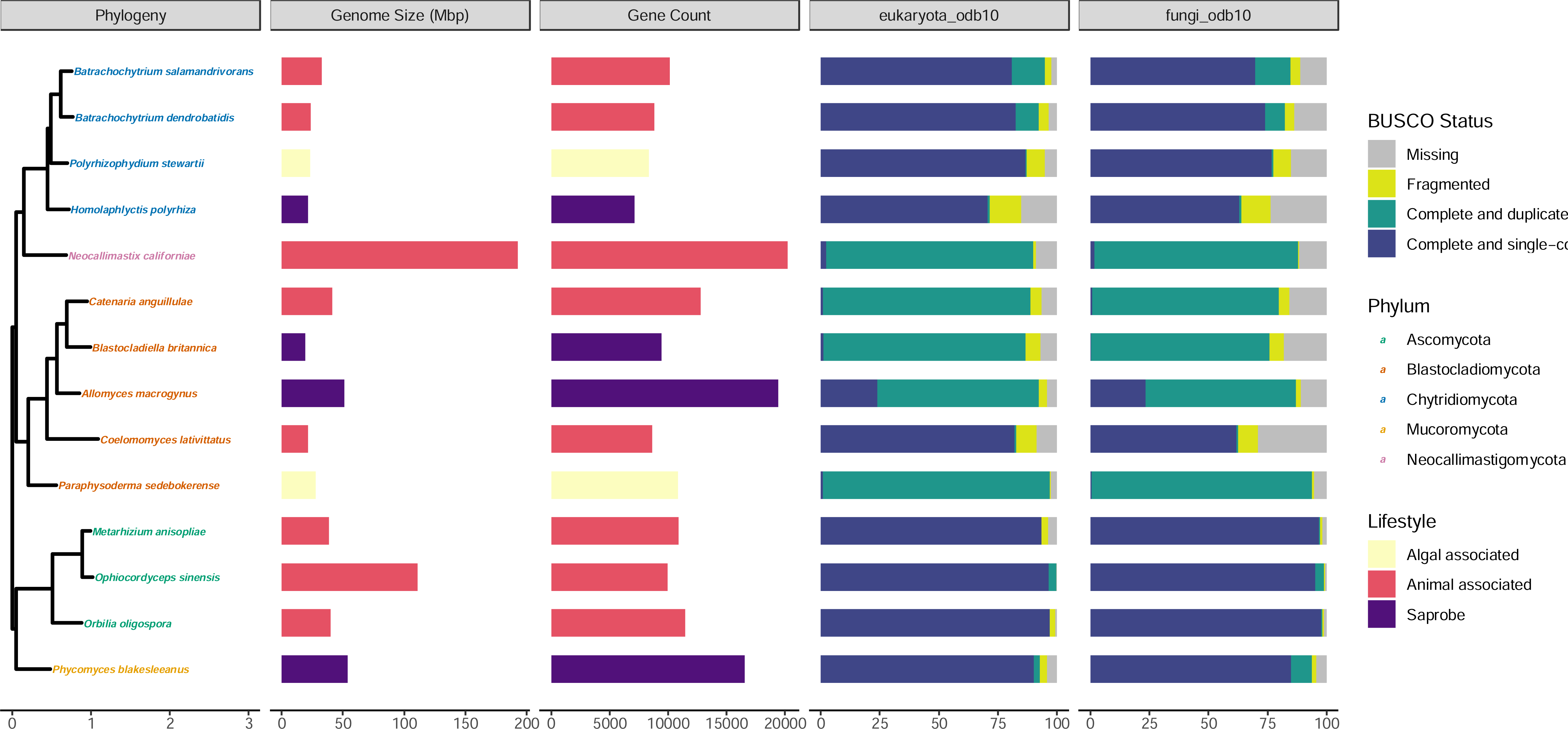
*C. lativittatus* protein set is comparable to those of other fungal taxa. Moving from left to right, here we show a maximum likelihood phylogeny which shows the relationship of *C. lativittatus* to other fungal lineages. This tree was generated using IQ-TREE2 on an alignment of BUSCO fungi_odb10 HMMs constructed using the PHYling_unified pipeline. Taxa labels in the phylogeny are shown colored by assigned fungal phylum. In association with this phylogeny, we then depict a barchart of the draft genome size (Mbp) for each taxon with genome size colored by fungal lifestyle (saprobe = purple, algal associated = yellow, animal associated = pink), followed by a barchart of predicted gene counts for each taxon with counts colored by fungal lifestyle. Next, we show barcharts of BUSCO ‘protein’ completion status for the eukaryota_odb10 and fungal_odb10 sets. Bars show the percent of genes found in each genome annotation as a percentage of the total gene set and are colored by BUSCO status (missing = grey, fragmented = yellow, complete and duplicated = green, complete and single-copy = blue). The values depicted here for *C. lativittatus* gene counts and BUSCO scores are based on the combined clustered transcript set (AOM90), and the genome size is the average size across all three assemblies. The BUSCO scores for individual *C. lativittatus* assemblies can be seen in Figure S2.

### Differential expression analysis reveals distinct expression profiles between life stages

Initial analysis of the transcriptome profiles supported a distinct divide between infection and sporangial life stages with each stage clustering separately (Figure 3A) and clear differences in expression in the most abundant differentially expressed transcripts (Figure 3B). We followed this analysis with differential transcript expression analysis using DESeq2 to identify specific transcripts responsible for these patterns.

**Figure 3.**
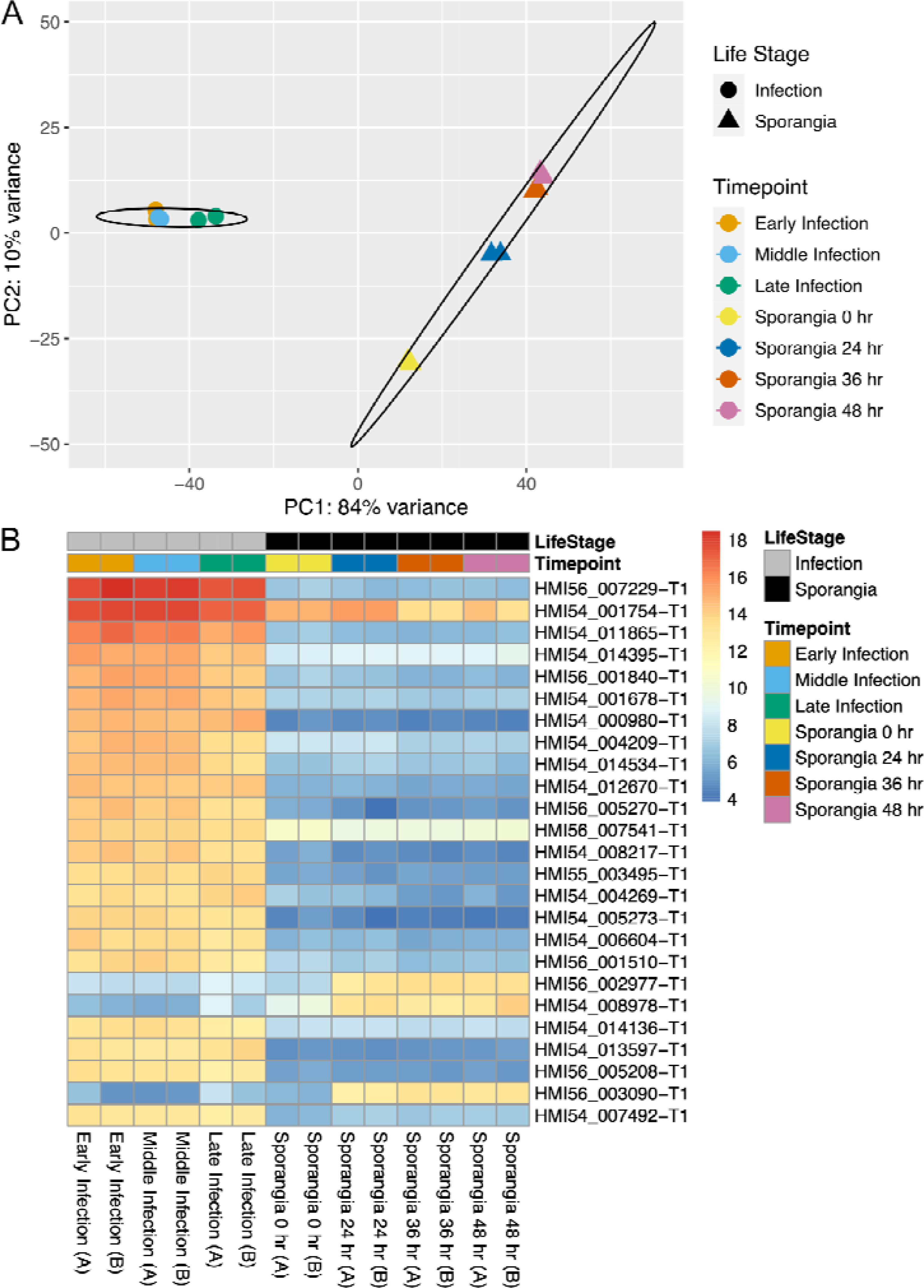
Transcript expression differs between *C. lativittatus* life stages. (A) Principal Component Analysis (PCA) of variance stabilized transcriptomic count data. Samples are colored by time points while shapes are used to broadly represent life stages (circle = infection, triangle = sporangial). Ellipses represent the 95% confidence interval around the centroid of each life stage. Replicate samples may be overlapping. (B) Heatmap showing the variance stabilized counts of the 25 most expressed transcripts with differential expression across life stages. Replicates are indicated by ‘A’ or ‘B’.

We found 1262 differentially expressed transcripts between life stages, with 395 transcripts enriched during infection compared to 867 enriched during sporangial life stages (*p* < 0.01, log2foldchange > 2). Of these, 575 (45.6%) had no matches to any of the databases used for annotation. Interestingly, while more transcripts were enriched in sporangial stages, many of the most abundant transcripts were representative of transcripts enriched in the infection stages and many of these transcripts were unannotated. For example, of the top 25 most abundant differentially expressed transcripts, 22 were upregulated during infection relative to sporangial stages (Figure 3B). Further, 18 of these 25 transcripts had no significant similarity to any features in the annotation databases, and further only one transcript was fully annotated, HMI54_014395 (*ERG10*), an acetyl-CoA C-acetyltransferase.

We performed GO enrichment analysis on the differentially expressed transcripts to identify enriched GO terms of interest (Table S3, *p* < 0.05). For the infection stages, GO terms from 38 biological processes (BP), 9 cellular compartments (CC), and 46 molecular functions (MF) were identified. Of these, seven of the top ten significantly enriched BP were metabolic processes (e.g., GO:0006082: organic acid metabolic process, GO:0046394: carboxylic acid biosynthetic process) and three were transport-related (e.g., GO:0006848: pyruvate transport, GO:1905039: carboxylic acid transmembrane transport). For the sporangial stages, GO terms from 35 BP, 26 CC, and 41 MF were identified. Of these, the most significantly enriched BP was related to reproduction (GO:0000003: reproduction), with four of the top ten significantly enriched biological processes related to metabolic and biosynthetic processes (e.g., GO:0005975: carbohydrate metabolic process, GO:0006183: GTP biosynthetic process, GO:0006228: UTP biosynthetic process) and three related to movement or organization of cellular machinery (e.g., GO:0006928: movement of cell or subcellular component, GO:0007010: cytoskeleton organization, GO:0007017: microtubule-based process).

### Differential expression analysis reveals complex pattern of expression clusters within life stages

Identification of differentially expressed transcripts across the development time course within sporangial and infection stages was done using a Likelihood Ratio Test (LRT). We found 380 transcripts that were significantly differentially expressed between infection stages (*p* < 0.01). Of these, we found two clusters of differentially expressed transcripts with similar expression patterns in each, group 1 and group 2 with 167 and 213 transcripts, respectively (Figure 4A). The same analysis with the sporangial stages indicated 3701 differentially expressed transcripts (*p* < 0.01) and out of which, we identified 7 clusters, each including differentially expressed transcripts with similar patterns of expression. There were 1083, 965, 859, 219, 355, 156 and 64 genes in groups 1 to 7, respectively (Figure 4B).

**Figure 4.**
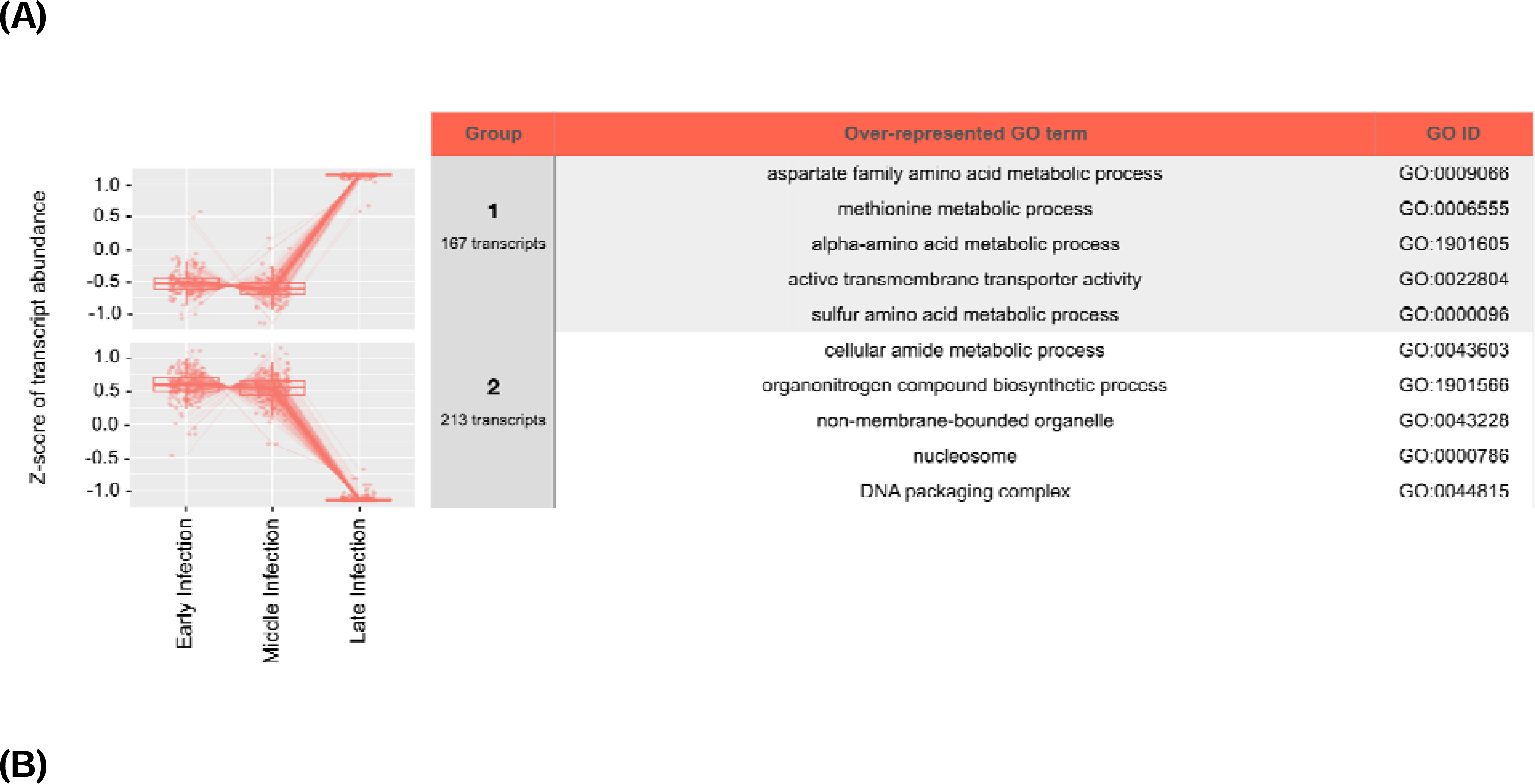
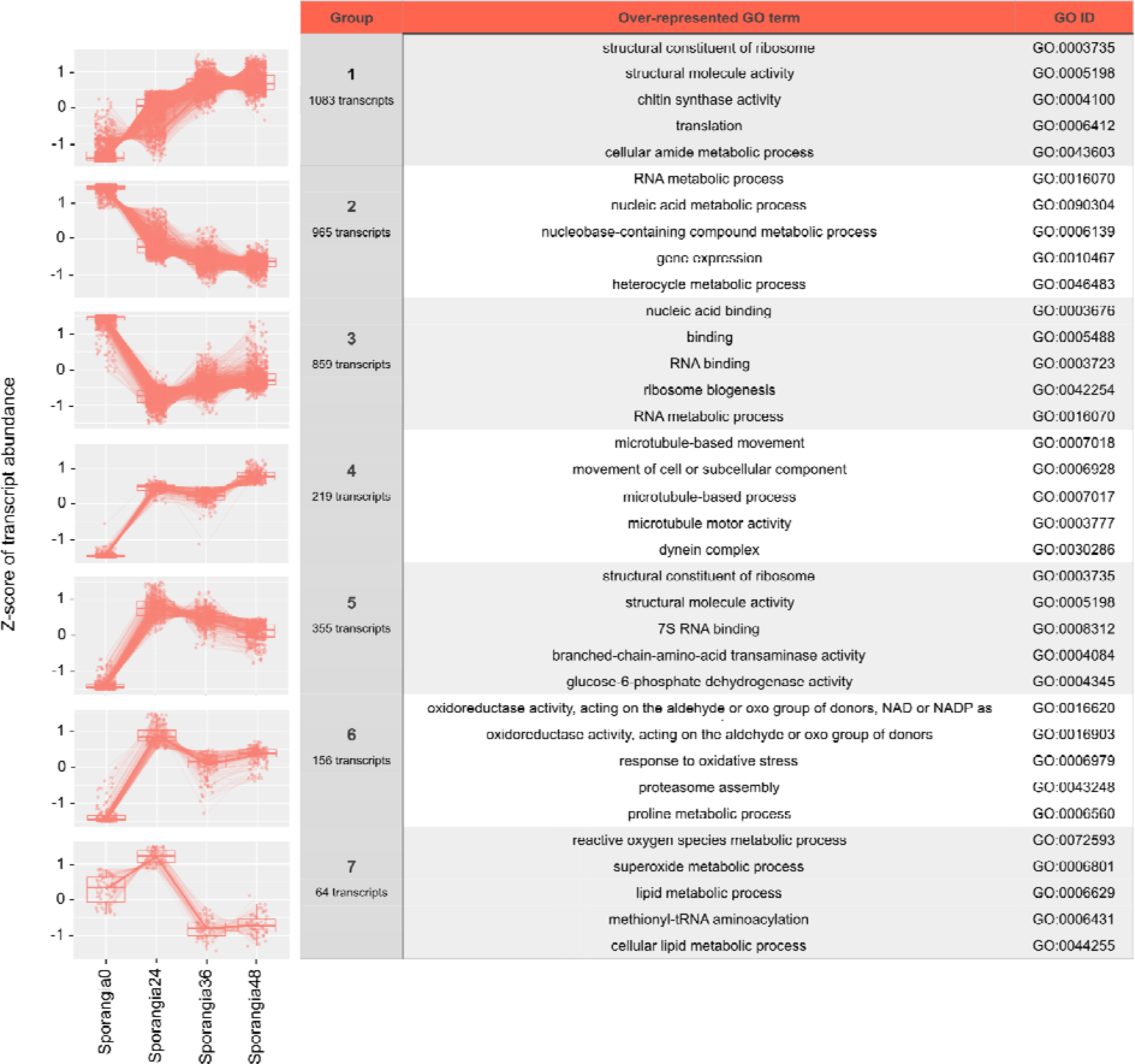
Transcript expression differs across the development time course conditions in each sporangium and infection stage. (A) The plots on the left are the two possible groups with specific transcript expression patterns across early, middle, and late infection timepoints and number of transcripts in each group. The table on the right shows the top five enriched GO terms for each of the groups of transcripts. (B) The plots on the left are the seven groups with specific transcript expression patterns across 0 hr, 24 hr, 36 hr and 48 hr timepoints within the sporangial life stages. The table on the right shows the top five enriched GO terms for each of the seven groups of transcripts.

In order to identify enriched GO terms of interest, we performed GO enrichment analysis on the differentially expressed transcripts across the developmental conditions within infection and sporangial stages (Figure S3, *p* < 0.05). Within the infection stages, we found 30 BP, 12 CC, and 33 MF enriched GO terms. Of these, the top significantly enriched GO terms were in the CC category (GO:0000786: nucleosome, GO:0043228: non-membrane-bounded organelle, GO:0043232: intracellular non-membrane-bounded organelle, GO:0044815: DNA packaging complex, GO:0032993: protein-DNA complex), and were also seen in group 2 with decreasing expression across infection stages (Figure 4A).

Furthermore, we found 19 BP, 10 CC and 28 MF enriched GO terms represented within the sporangial stages and of them, the most significantly enriched GO terms were related to structural molecule activity and binding (GO:0005488: binding, GO:0003735: structural constituent of ribosome, GO:0004100: chitin synthase activity, GO:0005198: structural molecule activity, GO:0003779: actin binding) as well as the GO terms related to peptide biological processes (GO:0006412: translation, GO:0043043: peptide biosynthetic process, GO:0006518: peptide metabolic process). Terms related to structural molecule activity and binding were generally seen in expression pattern groups 1 and 5 both of which generally increased in expression over time (Figure 4B).

### HMG-box loci were identified with differential expression across life stages

A total of five unique HMG-box genes were identified, with all five HMG-box genes present in the MEIOSPORE assembly and three in each of the AMBER and ORANGE assemblies. The identified HMG-box genes were found on small fragmented contigs (average: 8300 bp) which contained only a few neighboring genes (average: 3 genes). Despite their small size, synteny analysis across the three assemblies found that the AMBER and ORANGE assemblies share two HMG-box loci with each other. The third HMG-box genes in the ORANGE and AMBER assemblies were only shared with the MEIOSPORE assembly, which has four syntenic orthologous HMG-boxes (Figure S4). We tested the five HMG-box loci in *C. lativittatus* for synteny against *Allomyces macrogynus* to determine whether neighboring genes around HMG-boxes are conserved in sexually reproducing chytrids (Lee et al. 2010). We were unable to determine synteny of neighboring genes around HMG-box loci, possibly due to the fragmented scaffolds where these genes are found in our *C. lativittatus* assemblies. Phylogenetic analyses of the HMG-box genes showed that four of the HMG-box genes generally fell in clades with other blastoclads or chytrids, while HMI54_015288 fell into a clade with Dikarya. Further, three of the five HMG-box orthologs from *C. lativitattus* (HMI56_006544, HMI55_007199, and HMI54_004920) were present in a clade containing known mating-related HMG-boxes (Figure S5). HMI56_006544 and HMI54_004920 were present in all three *C. lativitattus* assemblies while HMI55_007199 was only present in the ORANGE assembly. Interestingly HMI56_006544 is closely related to SexM in *Phycomyces blakesleeanus,* however we were unable to identify a SexP homolog in *C. lativitattus.* The number of identified HMG-box genes here is in line with that of other blastoclads (i.e., *Blastocladiella britannica* and *Catenaria anguillulae*; Figure 5). Of the five identified HMG-box genes, HMI54_015288 was significantly over-expressed in the infection stages compared to the sporangial stages. (Figure S6; *p* < 0.01).

**Figure 5.**
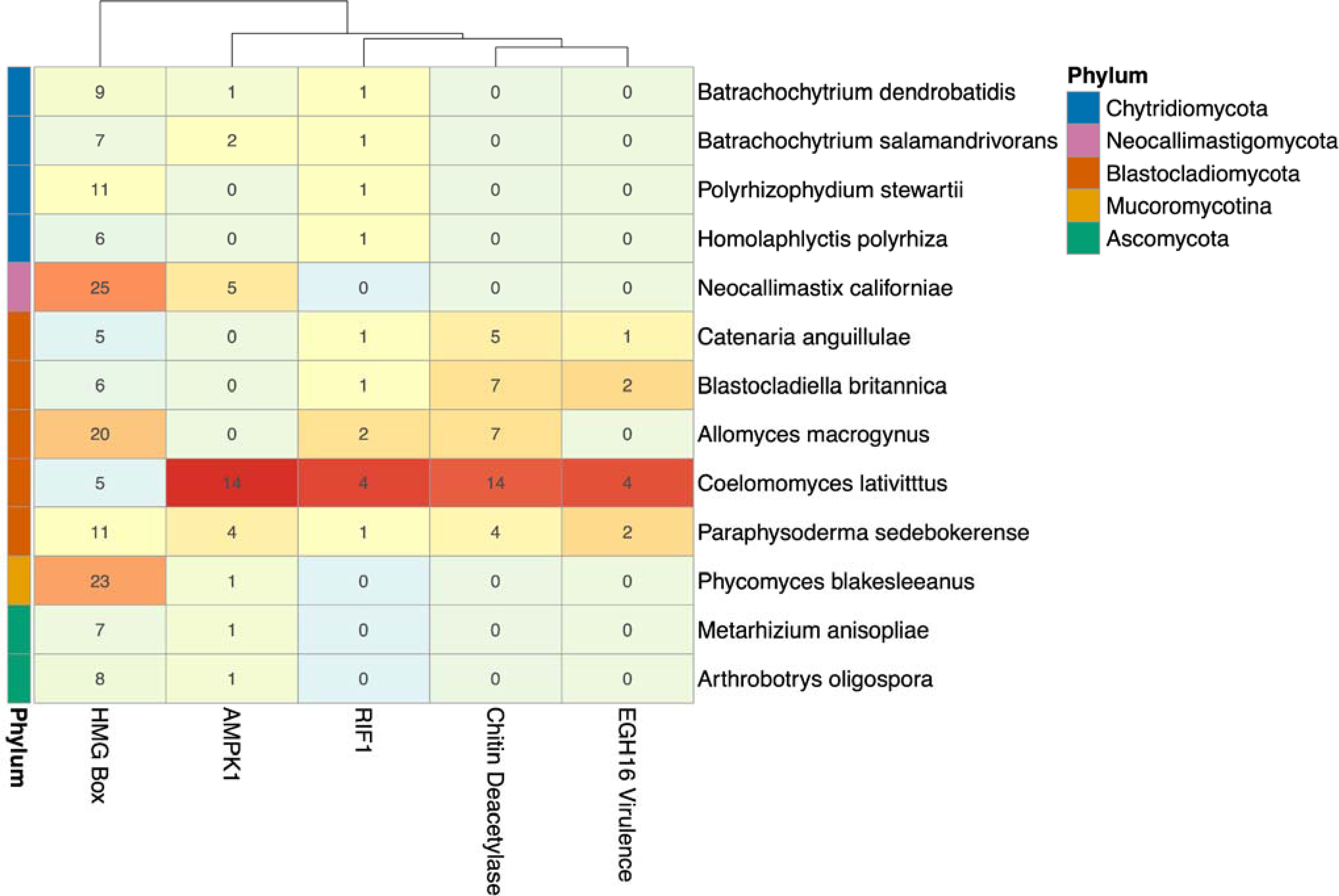
*C. lativittatus* copy number for HMG box, *RIF1*, chitin deacetylase, *Egh16-like*, and *AMPK1* gene orthogroups compared to other fungi. Here we depict a heatmap, organized by fungal phylogenetic relationships, depicting the copy number of orthologous genes, representing differentially expressed transcripts, with expanded gene counts in *C. lativittatus* relative to other fungi. The colors of the gene counts are normalized per gene family with red indicating a high normalized count and blue indicating a low count. The phylum for each taxon is indicated by the colored row labels at the left. The dendrogram above the heatmap clusters the columns by similarity in counts between the different gene families. Validated HMG boxes were found in two orthogroups which are shown combined here for simplicity.

*C. lativattatus may have expanded gene families of orthologs related to its unique life history* In order to determine if gene families expanded in *C. lativattatus* we performed Orthofinder analysis comparing *C. lativattatus* with other fungal taxa. In total we identified 37 398 orthogroups among the fungal taxa. Of these, 182 orthogroups were exclusively present in *C. lativattatus* and absent in all other fungal taxa.

We identified 10 orthogroups that both contained genes with differentially expressed transcripts and were expanded in copy number compared to other fungi. Of these 10 orthogroups, only four had predicted functional annotations (Figure 5). We tested the four orthogroups for duplication errors and removed any sequences that appeared truncated or had 100% sequence similarity to an ortholog from a different *C. lativittatus* assembly. Most of the copy number expansions for these four orthogroups appeared to occur on clades exclusive to *C. lativittatus.* The putative functions for the four validated orthogroups were replication timing regulatory factor 1 (*RIF1*), chitin deacetylase, adenosine monophosphate (AMP)-activated protein kinase 1 (*AMPK1*), and *Egh16*-like virulence factor. Within these orthogroups, only the *RIF1* orthogroup contained a gene that had higher expression in infection stages; the other three orthogroups had genes with higher expression during sporangial stages.

## DISCUSSION

### C. lativittatus annotated genomes are an important community resource

The genomes assembled and annotated here, while partial, are a promising and critical community resource as little genomic data exists for members of the Blastocladiomycota. The smallest public genome is *B. britannica* with a genome size of 19 Mb with 9431 predicted gene models and the largest is *Allomyces macrogynus* with a genome size of 47 Mb and 19 446 predicted genes (Grigoriev et al. 2014). *C. lativittatus* falls on the shorter end of this range with an average genome length of 21.5 Mb and average 7475 predicted gene models, possibly due to the incomplete nature of the draft genomes reported here. While partial, we think that the three *C. lativattaus* genomes assembled and annotated in this study provide much needed community resources for study of these obligate fungi. For example, the phylogenetic placement of Blastocladiomycota has been disputed, but inclusion of additional genomes like those reported here can help elucidate these ancient phylogenetic relationships (Amses et al. 2022).

### Transcriptomic landscape of C. lativittatus life stages provides insight into Blastocladiomycota biology

The transcriptome of *C. lativittatus* is a complex, dynamic, and underexplored landscape. The results of this study highlight a need for future refinement and exploration of gene annotation in this species, as evidenced by the 45.6% of differentially expressed transcripts with no annotation, and the majority of the top 25 abundant differentially expressed transcripts lacking functional annotation. In spite of these shortcomings, we were able to make generalizable insights about *C. lativittatus* biology from the subset of transcripts that were annotated with GO terms. Overall, during infection, GO terms were enriched related to metabolism and transport processes. While during sporangial stages, GO terms were enriched related to dispersal (i.e., cell signaling, locomotion and transport machinery). Looking at the expression patterns within life stages, we can begin to see more complicated trends emerge.

Within infection stages, we identified two different patterns of gene expression. In the first, gene expression increased over the course of development time (early, middle and late infection), with enriched GO terms related to membrane transport and metabolic processes. In the second, gene expression decreased over time with GO terms related to DNA replication, nucleic acid and amino acid biosynthetic processes.

The enrichment of metabolism and membrane transport processes in the transcriptomes during infection stages is similar to reports from other early diverging fungal lineages. Upregulation of transport-related pathways has been reported in chytrid infection of frog hosts, which the authors suggest might be related to nutrient availability and proliferation related to host association (Ellison et al. 2017). Further, in *Vavraia culicis* (Microsporidia) enriched GO terms for growth, metabolism and replication were identified and posited to be related to its generalist lifestyle and ability to infect multiple types of hosts (Desjardins et al. 2015). Here, we observed an enrichment of metabolism terms as part of a pattern of increasing expression (group 1, Figure 4A), and an enrichment of replication-related terms as part of a pattern of decreasing expression (group 2, Figure 4A). Thus, we may be observing a shift in priorities during infection, with early infection stages marked by increased replication as hyphagens grow into coenocytic hyphae inside the host and later infection stages marked by increased metabolism as the fungus proliferates and begins preparing to make sporangia.

Within sporangial stages, we identified seven expression patterns with two patterns displaying higher expression over these developmental stages, two patterns displaying lower expression, and three patterns with a relatively higher gene expression at the second time point (24 hrs), followed by stable or decreasing expression. In general, expression patterns with higher expression were enriched in GO terms related to chitin activity as well as terms related to dispersal and microtubule-based processes. In decreasing expression patterns the enriched GO terms were mostly related to metabolism and transcription. The other three expression patterns, which displayed the highest expression at the second developmental time point, were functionally different from each other. One group was enriched in GO terms related to dispersal and structural machinery, another in terms related to oxidative stress responses, and the third in terms related to lipid metabolism. Similarly, time series clustering of the transcriptome profiles of differentially expressed genes during the sporulation of the blastoclad, *Blastocladiella emersonii,* showed eight different patterns (Vieira and Gomes 2010).

The enrichment of reproduction and dispersal machinery-related mechanisms during the sporangial stages likely relates to the production of meiospores. For example, signal transduction and microtubule and cytoskeleton biogenesis were similarly reported to be enriched during sporulation in *B. emersonii (Vieira and Gomes 2010)*. These authors also observed a decrease in transcription and metabolism during sporulation which they attributed to the nutritional starvation required in order to sporulate. Additionally, previous investigations into protein synthesis in chytrids (Léjohn and Lovett 1966) and blastoclads (Lovett 1968; Schmoyer and Lovett 1969) suggest that translation does not occur until zoospore germination and that zoospores are likely partially dependent on maternal mRNA and ribosomes for initial protein production. Laundon *et. al*. (2022) posited that the chytrid zoospore life stage is optimized for dispersal to new hosts rather than general metabolism. The authors also reported complex lipid dynamics throughout the lifecycle of the chytrid, *Rhizoclosmatium globosum.* Of particular note, they observed increased expression of genes related to lipid transport and metabolism in zoospores, which often have large amounts of intracellular storage lipids. In this study across sporangial stages, we observed an enrichment of dispersal and microtubule-based terms as part of patterns with increasing expression, and transcription terms as part of patterns with decreasing expression (Figure 4B). We also observed one pattern of increasing and then decreasing expression related to lipid metabolism (group 7, Figure 4B). Therefore, here we may be discerning the metabolic preparation and production of meiospores for optimal dispersal, survival and host identification.

Later stages of host association are likely characterized by increased immune response in the host and countered by increased stress response by the fungus in order to continue to evade the host immune system. Upregulation of stress response genes has been reported in the plant pathogen *Zymoseptoria tritici* during late stages of sporulation in its host, which the authors posited might be protective (Keon et al. 2005). Similarly, we observed an enrichment of terms related to oxidative stress responses in the later stages of sporulation (group 6, Figure 4B), which we speculate may assist *Coelomomyces* evade host defenses during meiospore production.

### Mating-type genes may be useful for future work on evolution of sex in fungi

Unlike in animals and plants which have sex-specific chromosomes, sex determination in fungi is regulated by only a handful of genes. These mating-type (*MAT*) loci include HMG-box genes (Benkhali et al. 2013). While *MAT* loci in Dikarya have been widely studied (Wallen and Perlin 2018), the *MAT* loci of early diverging lineages of fungi have received relatively less attention (Idnurm et al. 2008). Given its obligate sexual two-host alternation of generations life cycle and the ability to separate sexed haploid gametes by color (orange or amber), *C. lativattatus* provides an intriguing system for investigating the evolution of sex in early diverging fungi. Using the genomes generated here, we identified five putative HMG-box genes, including one gene, HMI54_015288, which was differentially expressed between life stages, and three genes which were in a clade with known mating-related genes. Additionally, HMI56_006544, which was highly expressed during the sporangial stage, is homologous to the SexM gene of *Phycomyces blakesleeanus.* Interestingly, the three *C. lativitattus* HMG-boxes within the clade containing mating-related genes were up-regulated in the sporangial stage. Future work should tease apart the role in mating of these putative HMG-box genes in *C. lativattatus* and also place these genes in a comparative framework in order to further investigate the evolution of sex determination in fungi.

### Ortholog expansions in C. lativittatus may relate to host-association

Compared to other fungal lineages, *C. lativattatus* genomes had an enriched number of gene copies in four orthologs with functional annotations representing chitin deacetylase, *AMPK1*, *Egh16*-like virulence factor, and *RIF1* (Figure 5). We believe these expanded orthogroups may be good candidates for future investigations elucidating mechanisms behind *C. lativattatus*-host interactions.

In *C. lativattatus,* we found that members of the chitin deacetylase orthogroup were upregulated during sporangial stages and that “chitin synthase activity” was also an enriched GO term in group 1, which is a group represented by increasing expression (Figure 4B). Chitin deacetylases catalyze the deacetylation of chitin, an important structural component of fungal cell walls and insect cuticles, and have been previously reported in many fungal species (Zhao et al. 2010). Chitin-binding proteins and chitin deacetylation are thought to protect fungal pathogens against plant chitinases during infection (Gueddari et al. 2002; van den Burg et al. 2006), and have also been shown to be upregulated during infection of amphibian hosts by the chytrid pathogen, *Batrachochytrium dendrobatidis (Ellison et al. 2017)*. Further, the *B. dendrobatidis* genome has gene expansions of chitin-binding proteins which can confer protection against chintanse activity when expressed in the fungus, *Trichoderma reeseii (Abramyan and Stajich 2012; Liu and Stajich 2015)*. Thus, it is possible that the expansion in the chitin deacetylase orthogroup here may be related to *C. lativattatus* defense against its two hosts.

*AMPK1* genes are sensors that modulate energy metabolism and homeostasis, and can be important for regulating stress responses (Hardie et al. 2012). These genes can also be used to alter host energy metabolism by microbial pathogens (Mesquita et al. 2016). Increased counts of *AMPK1* orthologs and higher expression in sporangial stages in *C. lativattatus* may be related to regulation of increased stress during its two-host life cycle or to evasion of host immune detection.

In *C. lativattatus,* we found that members of the *Egh16*-like expanded orthogroup displayed higher expression during sporangial stages. *Egh16*-like virulence factors are proteins related to appressorium formation and pathogenesis which are present in pathogenic fungi including insect pathogens such as *Metarhizium acridum (Grell et al. 2003; Cao et al. 2012)*. *Egh16*-like factors have been postulated to aid in the penetration of insect cuticles (Cao et al. 2012). Thus, it is possible that the expanded *Egh16*-like virulence factor orthogroup is contributing to *C. lativattatus* virulence.

*RIF1* is important for telomere length control and subtelomeric silencing in fungi and other eukaryotes (Sreesankar et al. 2012). Subtelomeric regions have increased variability, caused by duplications and rearrangements that can result in functional novelty, including secondary metabolites related to pathogenicity and virulence. Silencing of subtelomeric regions is one way pathogens can control secondary metabolite expression (Wyatt et al. 2020; Diotti et al. 2021). Given the increased expression of members of the expanded *RIF1* ortholog during infection, these genes may have roles in silencing subtelomeric regions and may be another tool in the *C. lativattatus* toolbox for interacting with its hosts.

## CONCLUSION

We generated three draft genomes and annotations for *C. lativittatus* and characterized the *C. lativittatus* transcriptome landscape across infection and sporangial life stages. Little is known about the genomic landscapes of blastoclads or zoosporic fungi and thus, the genomic and transcriptomic data described here represent a valuable foundational resource. In the transcriptome investigation, we identified differentially expressed transcripts and enriched GO terms that provide insight into the blastoclad life cycle, with infection stages marked by an enrichment of metabolism and transport processes and sporangial stages by dispersal-related processes. Further, *C. lativittatus* has an interesting obligate alternation of generations life cycle with two hosts and here we found several ortholog expansions in virulence related genes which may have roles in modulating its host-associated lifecycle. As additional genomic data from other blastoclads and zoosporic fungi are generated, more powerful comparative approaches can be used to assess the evolutionary relationships of these lineages to other fungi, as well as better understand their complex life histories. We hope that this work sets the stage for this future studies by providing some foundational knowledge of these unique fungi.

## DATA AVAILABILITY

The decontaminated *C. lativittatus* genome assemblies and annotation are deposited at DDBJ/ENA/GenBank under the accessions JADGJU000000000, JADGJV000000000 and JADHYY000000000. The versions described in this paper are JADGJU010000000, JADGJV010000000 and JADHYY010000000. The raw sequence reads for the genome assemblies are available through BioProjects <u>PRJNA631428</u>, <u>PRJNA631429</u>, and <u>PRJNA631430</u>. The raw sequence reads for the transcriptome work are available under GEO accession <u>GSE222209</u>. Analysis scripts for this work are available on GitHub (https://github.com/stajichlab/Chytrid_Coelomomyces_Project) and archived in Zenodo (https://doi.org/10.5281/zenodo.7435008).

## ACKNOWLEDGEMENTS

Funding to support sequencing work was provided by a seed grant from the UC Riverside Office of Research and Chancellor’s Strategic Investment Funds project “*Coelomomyces Genomics for Mosquito Vector Control*”. CLE was supported by funds from California Department of Food and Agriculture agreement # 20–0267. MY was supported by funds from Gordon and Betty Moore Foundation (award #9337). J.E.S. is a CIFAR Fellow in the program Fungal Kingdom: Threats and Opportunities and partially supported by USDA Agriculture Experimental Station at the University of California, Riverside and NIFA Hatch projects CA-R-PPA-5062-H. Data analyses performed at the High-Performance Computing Cluster at the University of California Riverside in the Institute of Integrative Genome Biology were supported by NSF grant DBI-1429826 and NIH grant S10-OD016290. We thank Jericho Ortañez for assistance in experiments for generating genomic DNA and RNA.

