## Supplemental Table Legends and Figures for "Genomes and transcriptomes help unravel the complex life cycle of the blastoclad fungus, *Coelomomyces lativittatus,* an obligate parasite of mosquitoes and microcrustaceans"

**Table S1. Taxa utilized throughout this work.** We report the accession numbers, citations, and phyla for each reference draft genome used in this manuscript, as well as for the draft assemblies and combined transcript set generated here for *C. lativittatus.*

**Table S2. BUSCO scores.** We report the BUSCO scores for each genome and annotation, and the combined annotation gene set (AOM90) used for transcriptome analyses. For comparison, we provide BUSCO scores as well for genomes and annotations of Blastocladiomycota and other diverse fungal lineages. BUSCO scores are reported using ‘genome’, ‘transcriptome’ and ‘protein’ modes for the eukaryota_odb10 and fungi_odb10 sets.

**Table S3. Enriched GO terms for genes that are differentially expressed between life stages.** Enrichment analysis identified GO terms that were significantly enriched in the infection and sporangial stages (*p* < 0.05). These terms were summarized and filtered with Revigo. Here we provide the GO ID, GO term, GO category (BP: biological process, CC: cellular component, MF: molecular function) and *p*-value for each enriched term.

**Figure S1. Assessment of microbial contamination in the *C. lativittatus* str. AMBER, ORANGE and MEIOSPORE assemblies.** (A) Visualization from the anvi’o pipeline for the AMBER assembly. The hierarchical clustering of contigs based on differential coverage and tetranucleotide frequency, followed by contig length and GC content are shown. Next, coverage of each genomic sample (AMBER, MEIOSPORE, and ORANGE) to the AMBER assembly is depicted. Finally, taxonomy per contig is colored by superkingdom (eukaryota = blue, bacteria = yellow) and the possible contaminant contigs are labeled and highlighted in the hierarchical tree structure in yellow. BlobTools2 visualization of the (B) AMBER, (C) ORANGE, and (D) MEIOSPORE assemblies. Each circle represents a contig in the assembly, scaled by square-root normalized length, and colored by superkingdom (eukaryota = blue, bacteria = yellow). On the X axis is the average GC content per contig and on the Y axis is the average coverage per contig. The marginal histograms show cumulative genome length (Mb) GC content bins (X axis) and for coverage (Y axis).

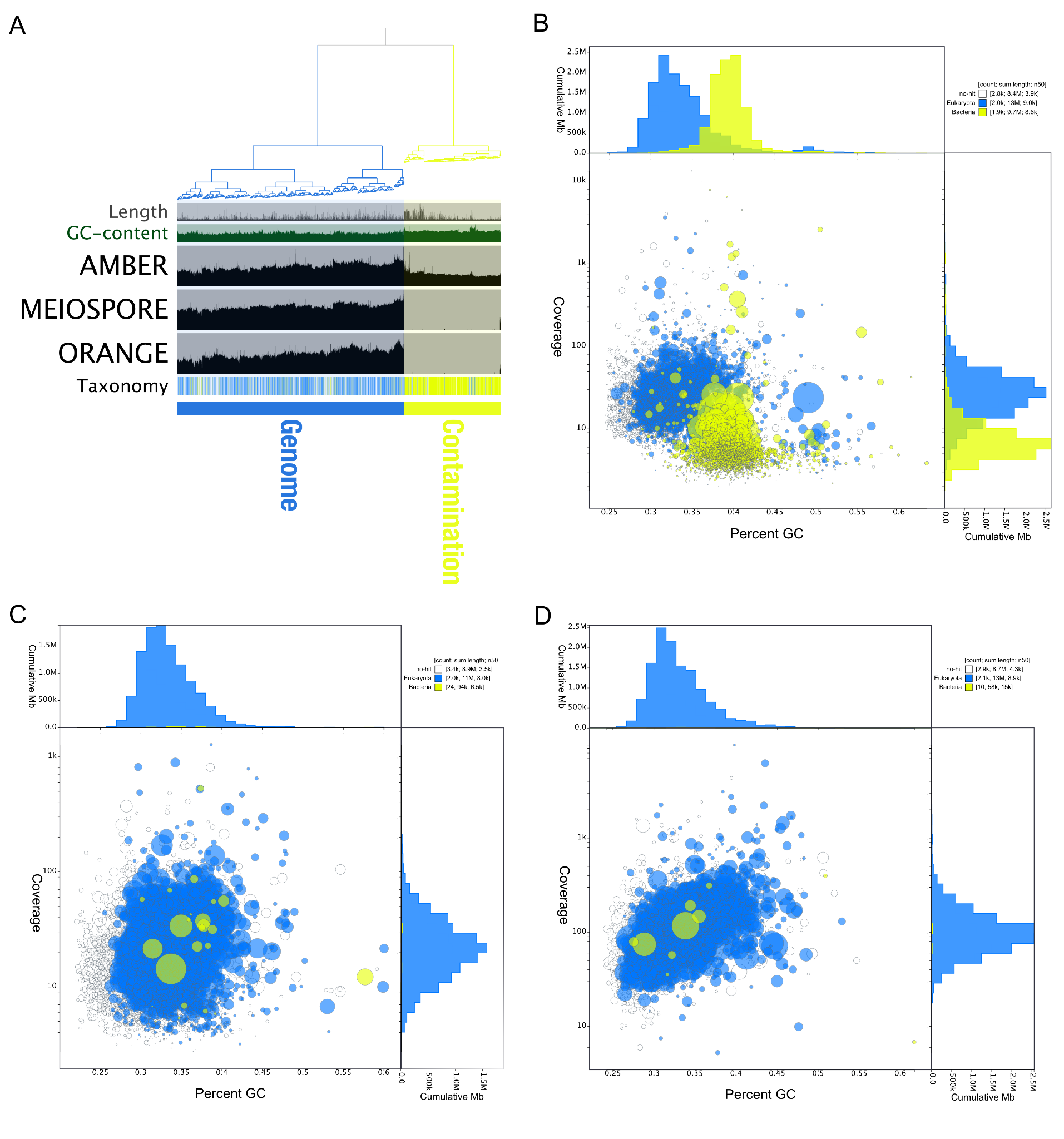

**Figure S2. Combined transcript set (AOM90) has improved BUSCO scores compared to individual gene sets.** BUSCO was run in ‘protein’ mode on individual assemblies and then the combined transcript set (AOM90). Here we show barcharts of the BUSCO score results for each of the eukaryota_odb10 and fungal_odb10 maker gene sets. Bars show the percent of genes found in each genome annotation as a percentage of the total gene set and are colored by BUSCO status (missing = grey, fragmented = yellow, complete and duplicated = green, complete and single-copy = blue). These values can also be found in Table S1.

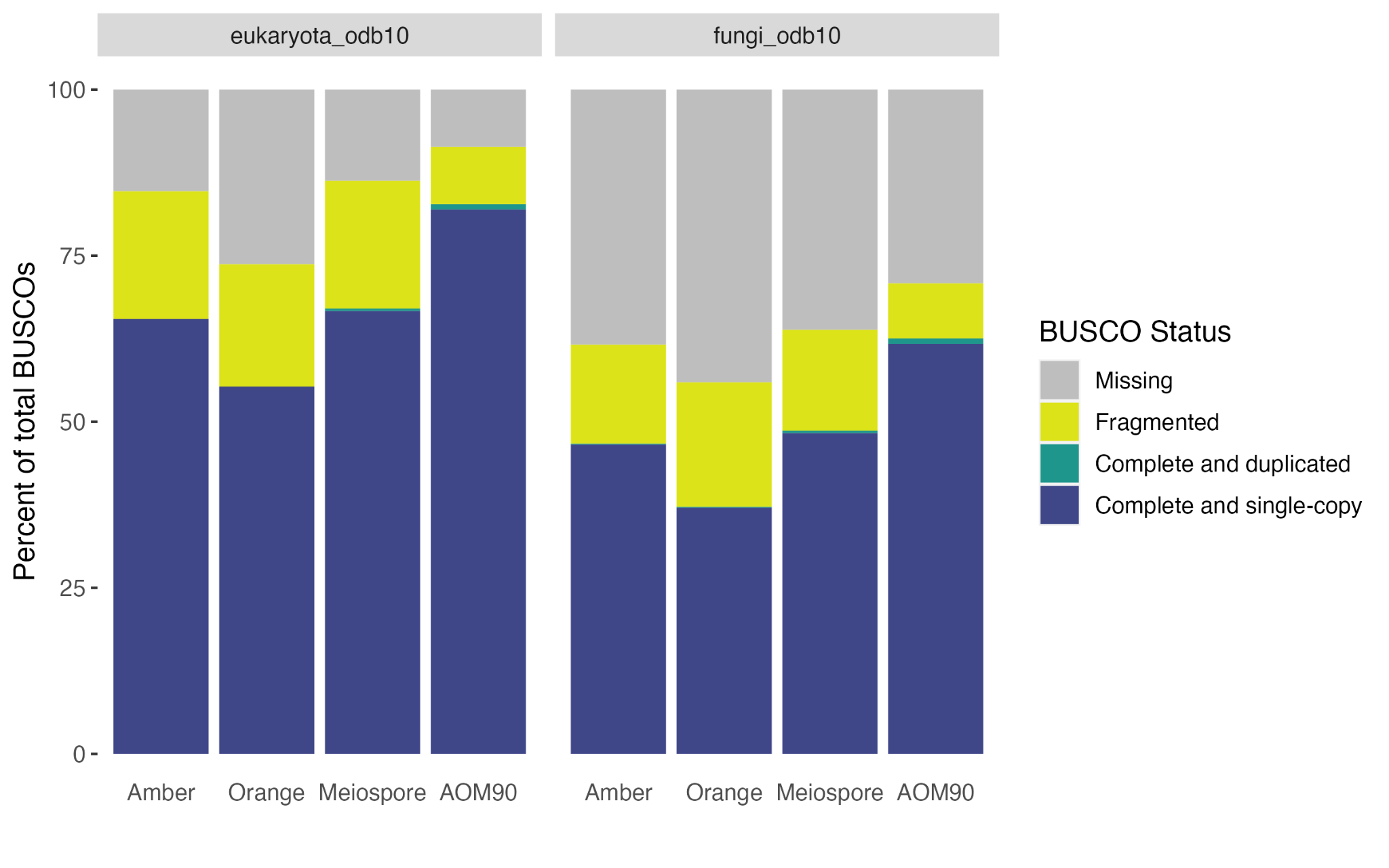

**Figure S3. Results of Gene Ontology enrichment analysis performed at each of three GO classes: biological processes (BP), molecular functions (MF), and cellular components (CC).** (A) Infection stage, (B) Sporangial stage. The Y-axis lists REVIGO filtered and non-redundant enriched GO terms and the X-axis and the bubble size indicate the number of genes associated with a GO term. Bubbles are colored by the p-value (FDR < 5%). Each GO class (BP, CC, MF) is represented by a different color (pink, yellow and purple, respectively).

**(A)**

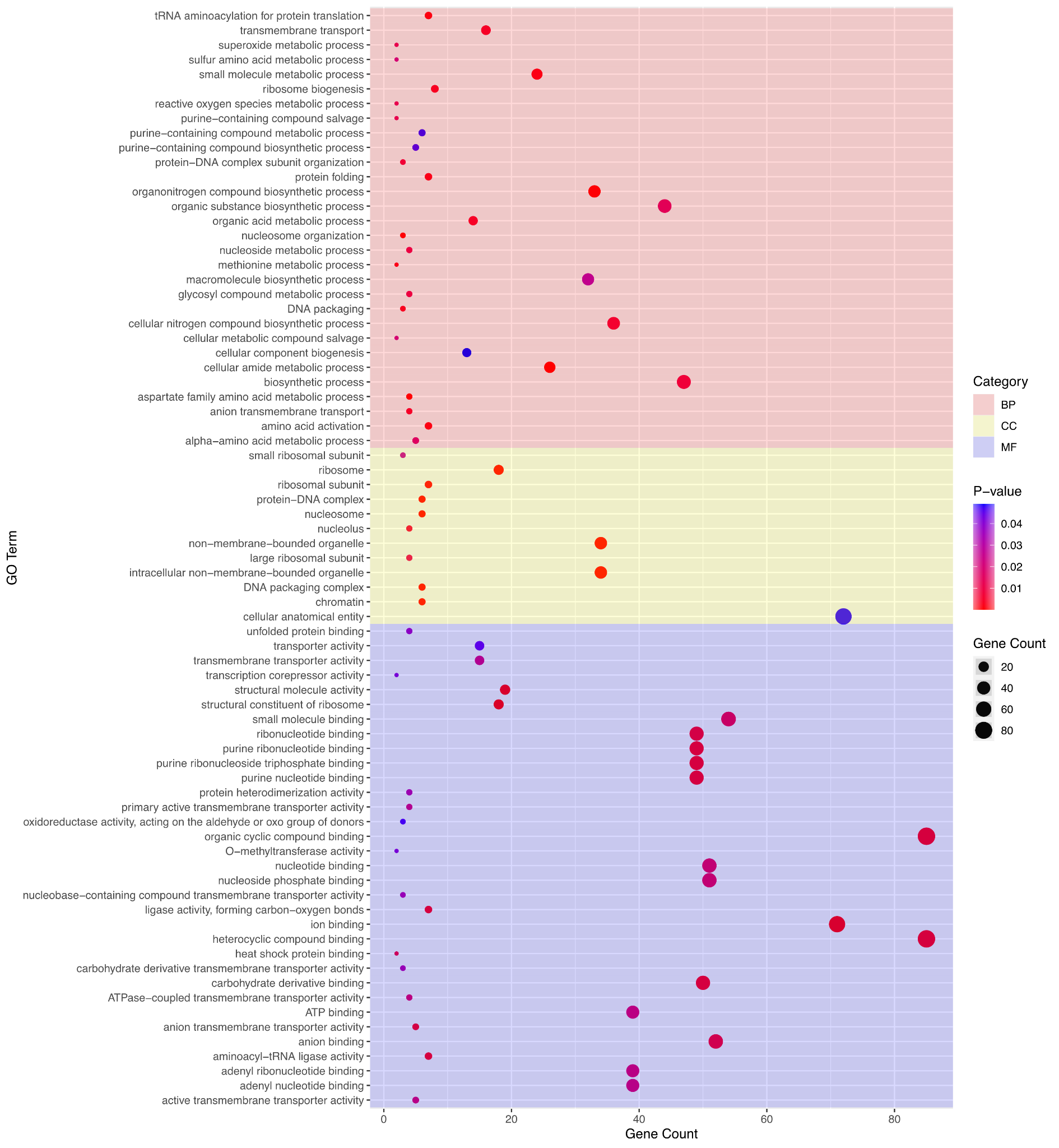

**(B)**

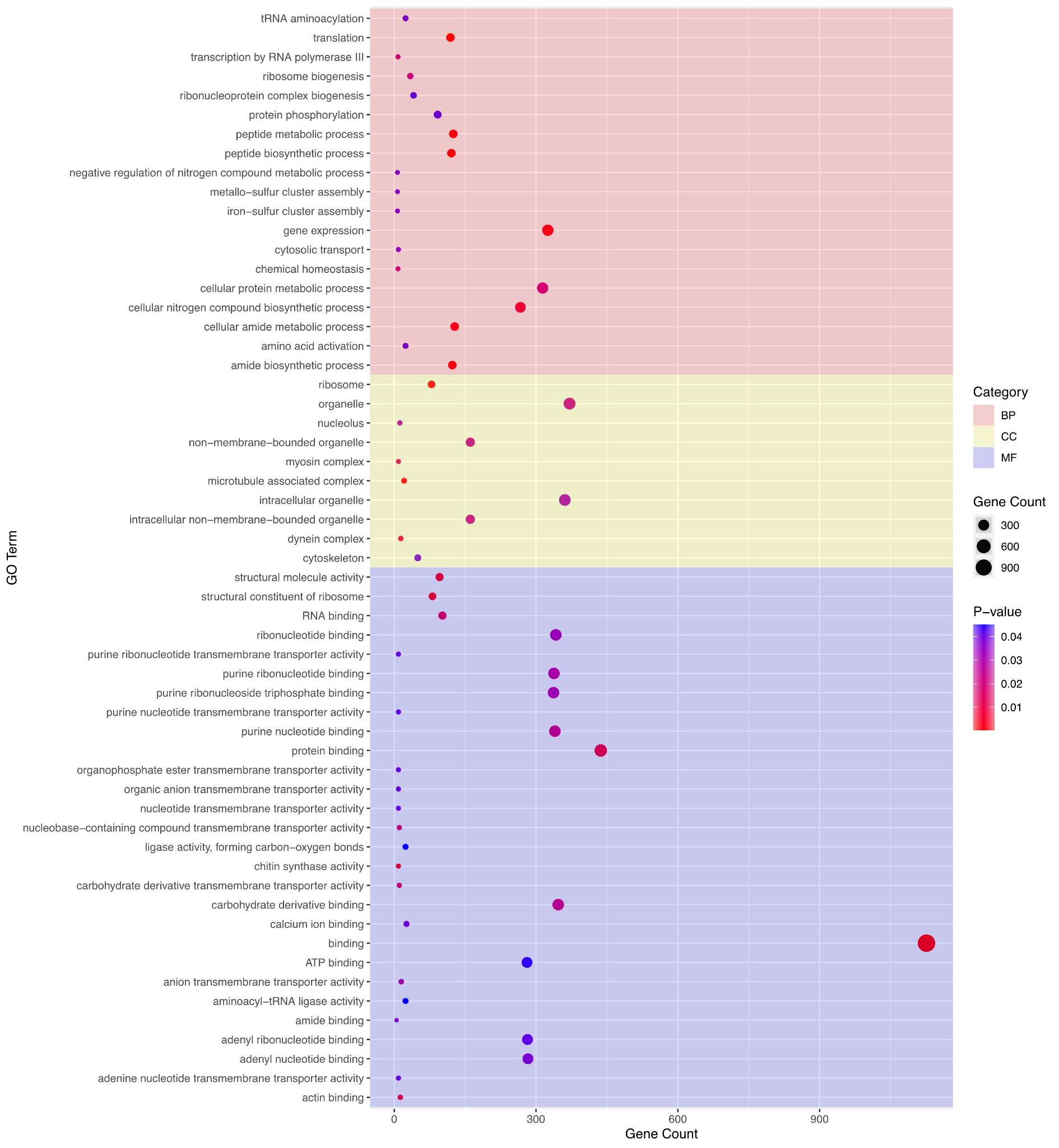

**Figure S4.** **Synteny of *Coelomomyces*** **MAT loci.** Here we show synteny of the five HMG box orthologs across the three *Coelomomyces* genome assemblies. The gray-scale color bar indicates percent amino acid sequence similarity between connected proteins. Colored arrows represent proteins with shared identity, while gray arrows had no orthology to HMG box regions.

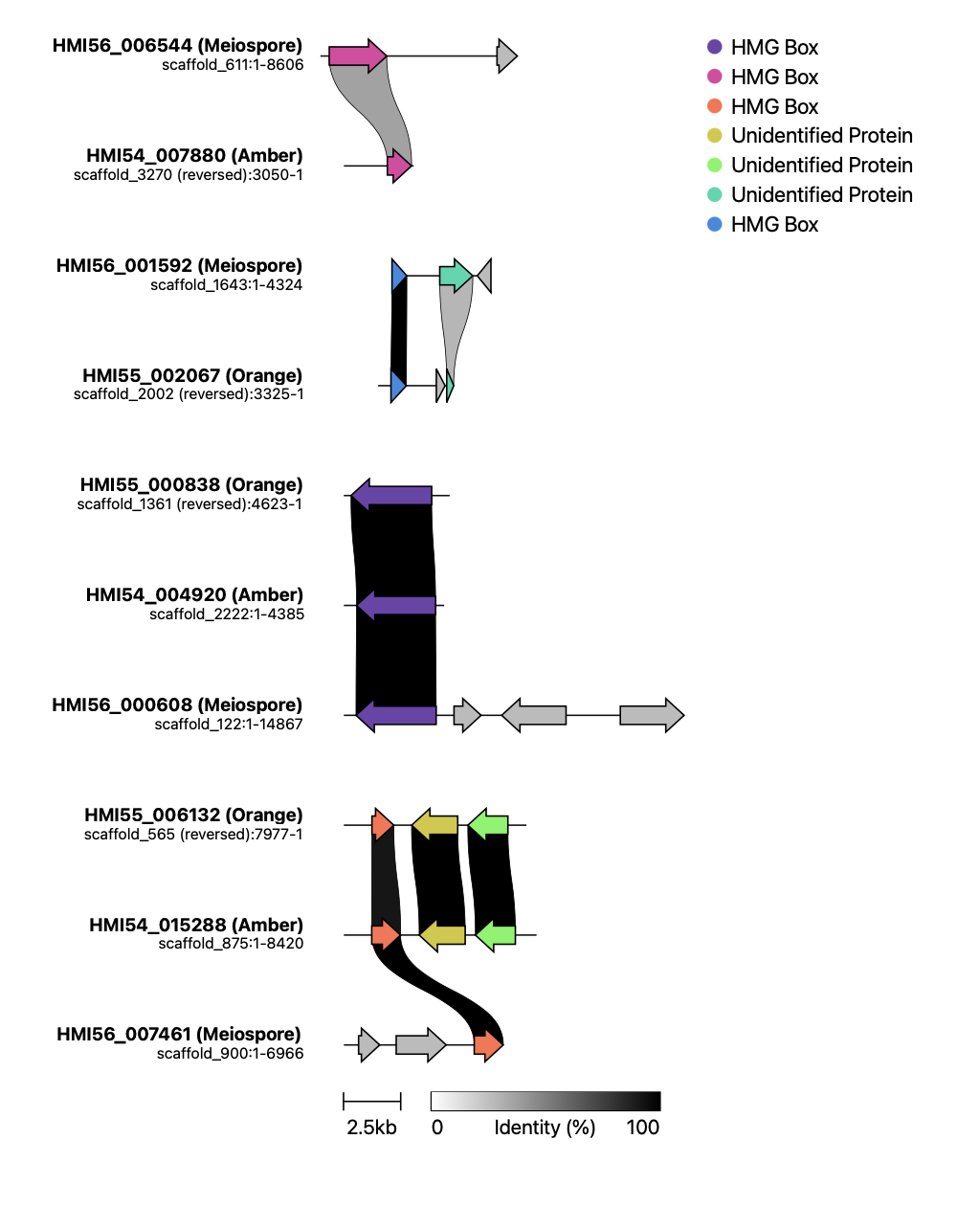

**Figure S5. Phylogenetic placement of *Coelomomyces*** **MAT loci.** Here we show the phylogenetic placement of the five HMG box genes from the MEIOSPORE assembly with HMG boxes from other fungal species. MAT loci in other fungi are indicated with labels indicating their gene name. Only HMG Box genes experimentally confirmed to be involved in mating are labeled by their gene ID in the colored boxes. Bootstrap values are represented by colors at each node and the fungal phylum for each gene is indicated at each tip by a colored circle. The clade of HMG orthologs without any members involved in mating has been collapsed.

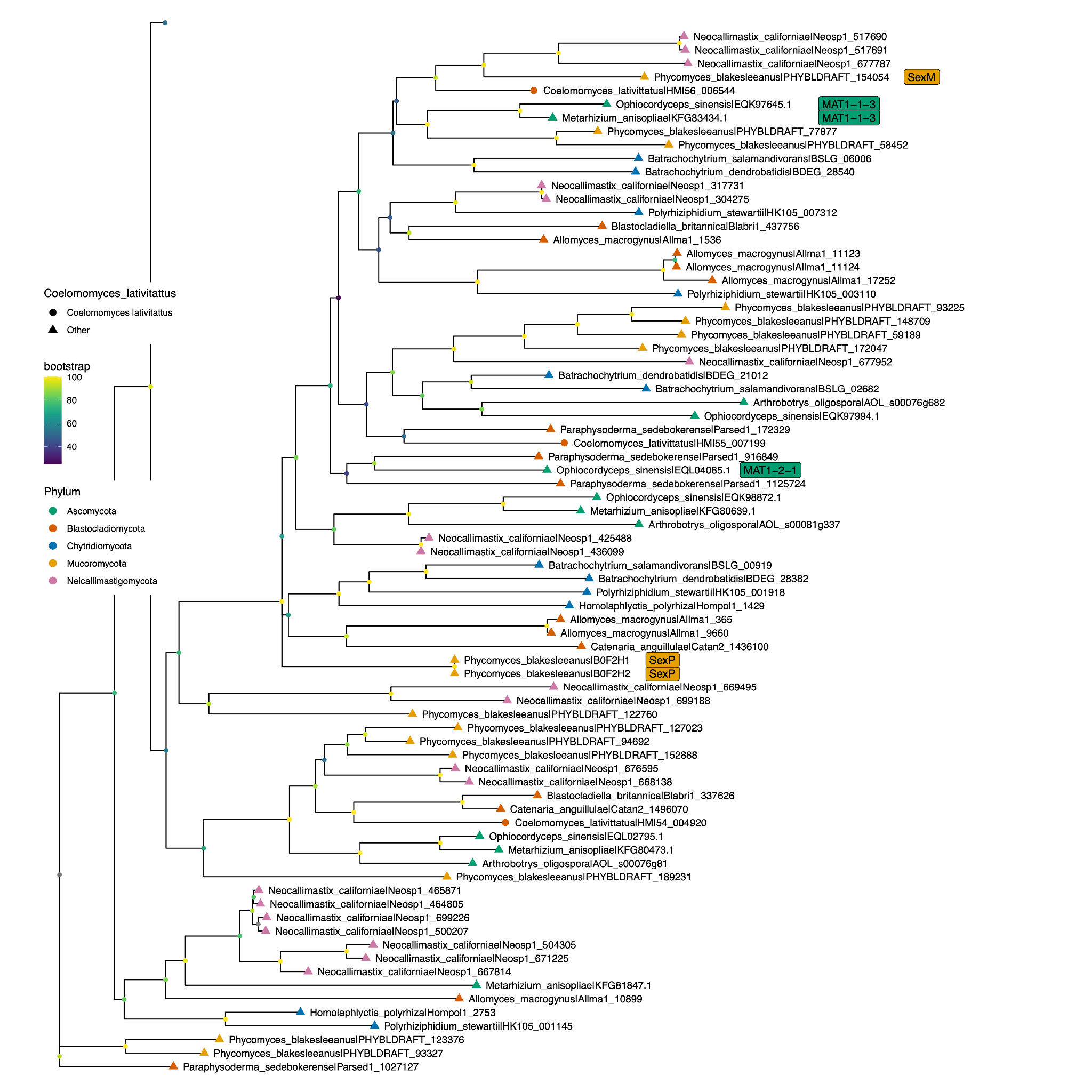

**Figure S6. MAT loci expression across life stages.** Here we depict variance stabilized transcriptomic counts of the five HMG box orthologs across *Coelomomyces* life stages (A-E). (B) HMI54_015288 is significantly upregulated during infection relative to the sporangial stages (*p =* 8.456e-06). (E) In contrast HMI55_007199 was significantly upregulated during the sporangial stage compared to infection stages (*p* = 2.27e-05). (A,C,D) The remaining HMG Box genes were not significantly differentially expressed between the life stages (*p* > 0.05).

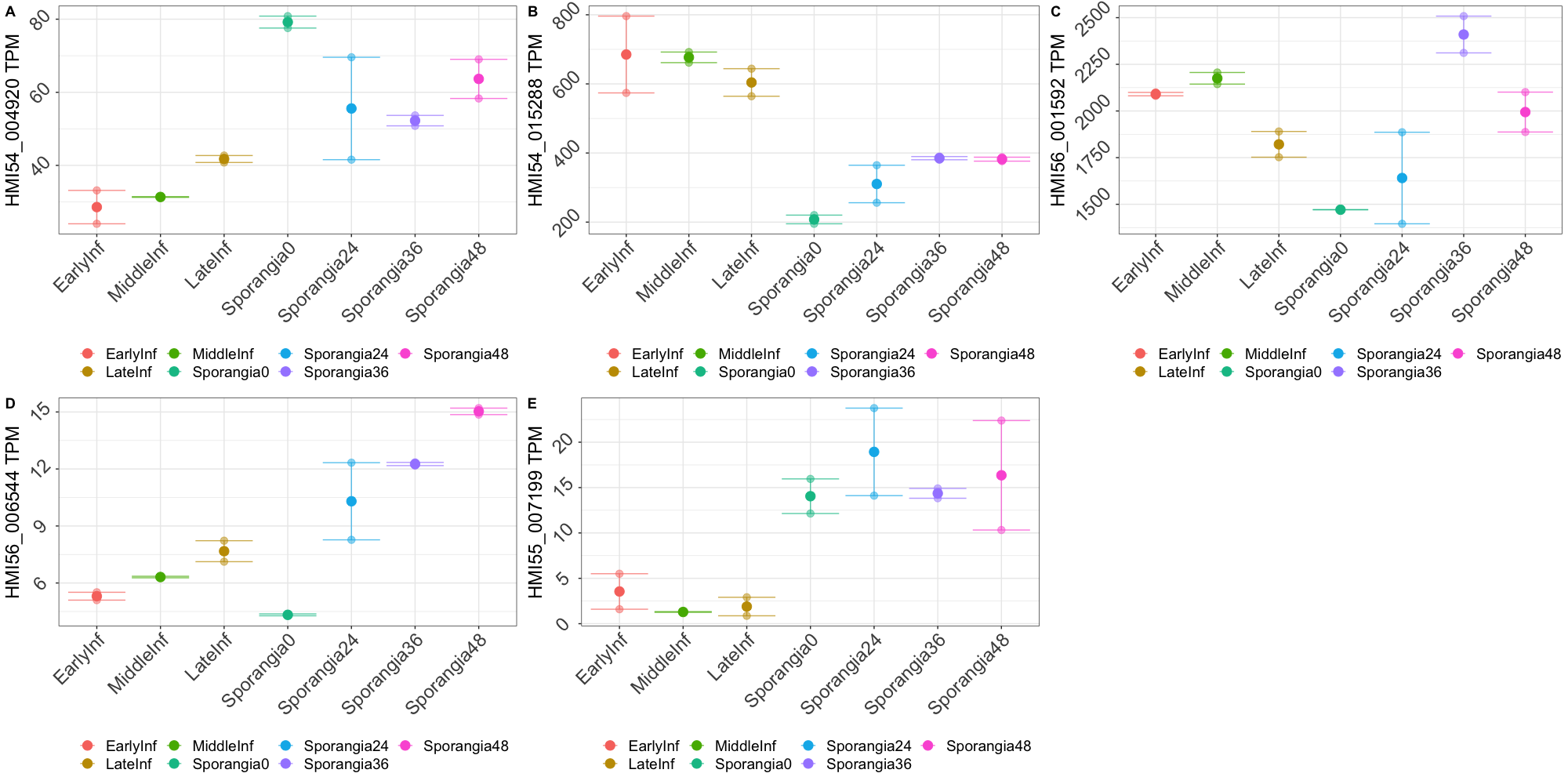
